# Kernelized approach enables explainable gene prioritizations for complex traits

**DOI:** 10.64898/2026.07.08.737338

**Authors:** Taotao Tan, Md. Abul Hassan Samee

## Abstract

Genome-wide association studies (GWAS) have identified numerous variant-trait associations; yet, assigning effector genes to GWAS loci remains challenging. Similarity-based machine-learning methods, such as PoPS, prioritize effector genes from shared functional profiles among trait-relevant genes. These models assign a prioritization score for each gene and nominate a single effector gene within a GWAS locus. However, the scores provide limited insight into why a gene was prioritized or whether the nomination is biologically plausible. To address this gap, we introduce Kernelized Polygenic Priority Score, K-PoPS, a kernelized reformulation of PoPS that enables gene-centric explanations by decomposing each prediction into contributions from training genes. For each prioritized gene, K-PoPS reports top contributor genes and an anchor score that quantifies support from a user-defined set of trait-relevant genes. Across 38 Pan-UK Biobank traits, the full-feature OLS implementation underlying K-PoPS improved closest-gene enrichment relative to default PoPS for 26 of 37 evaluable traits. Across 25 traits with curated anchor sets, predictions supported by anchor scores were more enriched for closest-gene proxies than unsupported predictions. When applying to blood level apolipoprotein B, K-PoPS nominated *SCARB1* over *UBC* gene, and further provided convincing explanations that support this prediction. Using explanation evidence, K-PoPS identified multiple plausible effector genes within a dilated cardiomyopathy locus, contrary to the parsimonious assumption. In summary, K-PoPS provides a post hoc framework for examining and interpreting GWAS effector-gene nominations.

## Introduction

Genome-wide association studies (GWAS) have uncovered thousands of loci associated with complex traits, yet mapping these loci to effector genes remains a major challenge^1^. The difficulty arises because significant variants often lie in non-coding regions or are in linkage disequilibrium (LD) with causal variants, overall making it unclear which gene(s) drive the association. This gap between statistical associations and biological insight led to the development of methods that integrate functional data to prioritize effector genes. Broadly, gene prioritization approaches fall into two categories: 1) locus-based methods that use local genomic information (e.g., nearest-gene, coding mutations, enhancer contacts, eQTLs) to assign effector genes to causal variants^2,3^, and 2) similarity-based methods (e.g., PoPS) that find effector genes across the entire genome with shared etiological functions, pathways, or network connections^4–7^. For instance, PoPS nominated novel genes such as *LGR4* for estimated glomerular filtration rate^6^. FLAMES prioritized genes in the *FSHB* locus related to dizygotic twinning^7^.

These computational approaches have significantly advanced post-GWAS analysis by streamlining effector gene prediction. However, the prioritized gene lists are not always biologically plausible, and different methods often produce inconsistent results. For example, a recent study found that only 50% – 75% of prioritized genes are reproducibly identified for the same disease^8^. When benchmarking between gene prioritization tools, *Weeks et al.* found that PoPS has 26% – 52% concordance with other methodologies^6^. We further examined PoPS-prioritized genes for 18 disease traits and found that only 0% – 33% of these gene-trait links have been previously reported in the Human Phenotype Ontology (HPO) database (see result section). These newly identified gene-trait links are not necessarily statistical artifacts, but these candidate genes should be interpreted cautiously. Relying solely on the gene prioritization scores for downstream experimental validation and therapeutic target selection can be of high risk in terms of both cost and time. Taken together, the community requires guardrails for gene prioritization programs to ensure transparent and interpretable predictions^8^.

Explainable AI (XAI) techniques have been developed to enhance model transparency, and were widely used in various applications such as medical imaging^9,10^. These techniques ensure that the predictions are inspectable, and humans can detect artifacts and make informed judgements. Most XAI tools, such as saliency maps and SHAP values^11^, can attribute predictions to input features. By examining these feature attributions, it enables answering questions such as “what regions in an X-ray contributes to the prediction of cancer?” Through explanations, clinicians can reason if a prediction is trustworthy and re-evaluate medical decisions. However, XAI has yet to be implemented in gene prioritization, a field where predictions guide expensive functional follow-up. One major obstacle is that genomic features are prohibitively high-dimensional and heterogeneous. For example, PoPS predicts effector genes using more than 55,000 features derived from RNA-seq, pathways and biological networks. Although pathways and networks are relatively interpretable, the majority of features (approximately 40,000 features) are derived from various RNA-seq analyses, such as Principal/Independent Component Analysis (PCA, ICA) loadings of a specific single-cell dataset. Even when applying those feature-centric XAI methods to PoPS, it may attribute the prediction to these hard-to-interpret features.

To address the interpretability issue for high-dimensional and heterogeneous gene features, we propose to apply an example-based (gene-centric, in this context) XAI framework on gene prioritization. Instead of attributing predictions to features, this class of approaches, such as influence functions^12^ and representor points^13^, pinpoint the most influential training genes for each prediction^14^. This framework shifts the question from “which features drive the prediction” to “which genes support the prediction”. The latter arguably offers greater biological interpretability, since genes are generally well-defined and well-studied units. Using the gene-centric explanations, researchers could evaluate if the prioritized genes are biologically plausible, generate new hypotheses, and thereby enable more informed decisions regarding downstream experimental validation.

## Results

### Overview of K-PoPS

PoPS first assembled high-dimensional gene-level features *X ∈ R^n^*^×57,543^ from 1) principal components (PC) and differential gene expression analysis for 77 bulk & single cell RNA-seq datasets; 2) pathway memberships from KEGG, GO, Reactome, and Mouse Genome database; 3) PPI network derived InWeb_IM network (Figure 1a). PoPS then calculates gene-level association statistics across all the genes with MAGMA: *y ∈ R^n^* (*n* genes in total). Using gene-level features *X ∈ R^n^*^×57,543^, PoPS then conducts feature selection, and predicts MAGMA scores from those features using ridge regression: *y* = *X* β. After obtaining feature weights 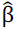, PoPS score 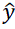 can be calculated as: 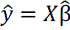. Under the leave-one-chromosome-out (LOCO) scheme, the learned coefficients 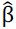 were determined using genes from the training chromosomes {*X*_train_, *y*_train_}, and then applied to the test chromosome 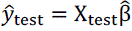. The process is repeated across all chromosomes, and the resulting scores are combined to generate the final PoPS score. After obtained the PoPS scores, one typically ranks all the genes within a locus, and select the gene with the highest PoPS score^6,8^.

**Figure 1.**
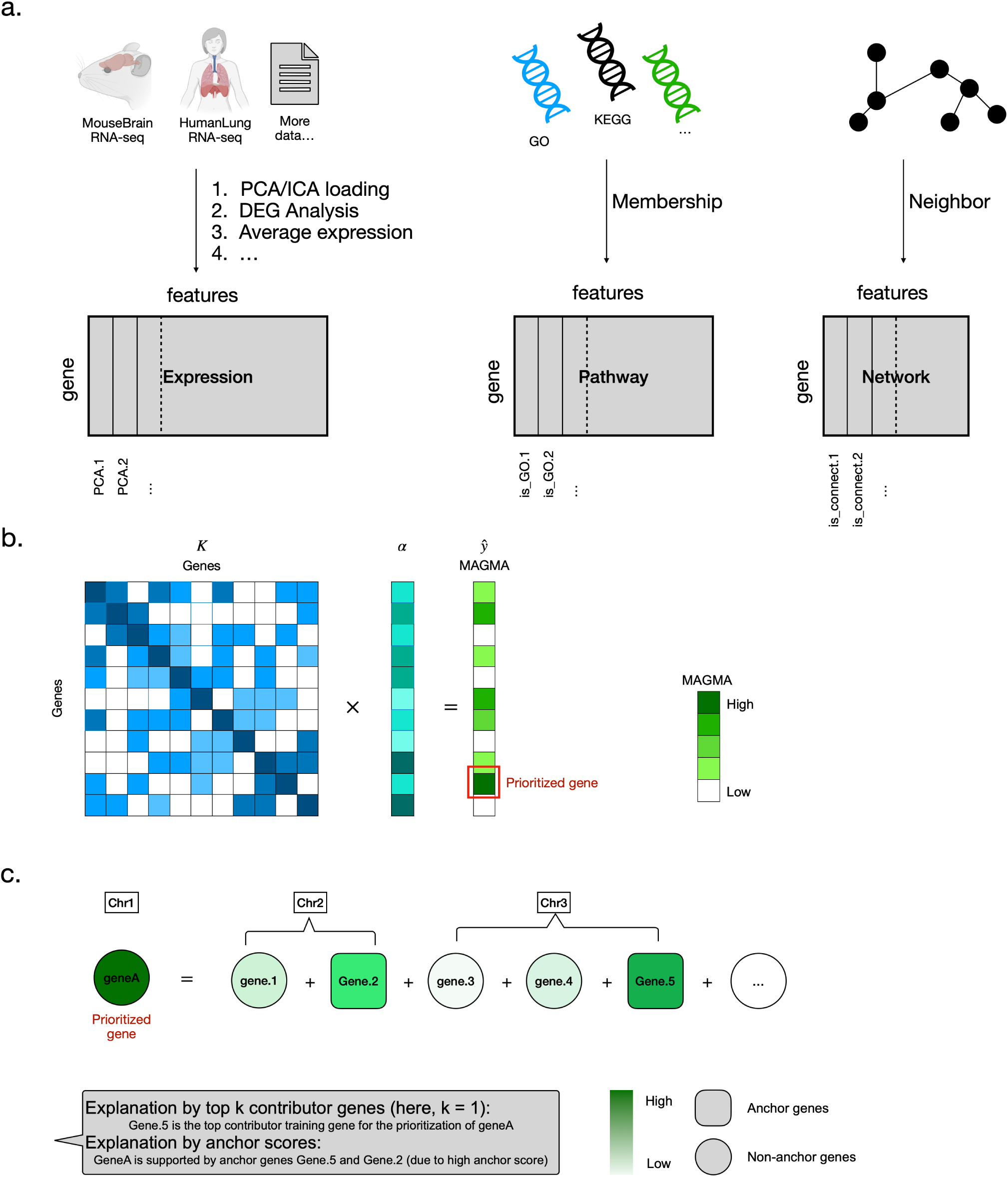
a) PoPS curated over 55,000 gene features from expression, pathway and networks. Expression features are derived from various analysis (PCA/ICA loadings, DEG analysis, etc) from 77 dataset. Pathway membership features are derived from 4 databases including GO, KEGG, Reactome, and Mouse Genome databases. Network features are derived from InWeb_IM PPI network. b) K-PoPS first constructs the kernel matrix which measures the similarity between genes. K-PoPS then predicts the MAGMA score from the kernel matrix. c) For the prioritized genes, K-PoPS assigns contributions to each training gene. It identifies the top contributor genes, and quantifies the support from pre-defined anchor genes.

We enabled gene-centric explanations by reformulating PoPS as a kernel regression. In computer vision tasks, kernel regression was first used by *Yeh et al.* to explain model decisions of deep neural network^13^. Specifically, kernel regression computes the gene prioritization score as a weighted sum of similarities between a given test gene and the training genes: 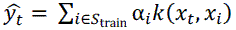. Here, *x_t_* is the feature vector for the test gene, *x_i_* is the feature vector for the training gene *i*, *S*_train_ is the gene set that includes all genes in the training chromosomes, *k* is the kernel function (similarity evaluation), 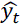 is the predicted value (i.e. the K-PoPS score) for the test gene^13^. This formulation effectively decomposes a gene prioritization score into the contributions of each training gene. Examining the contribution scores α*_i_k*(*x_t_*, *x_i_*) can help elucidate model decisions, as it pinpoints influential training genes for a given prediction^13^.

K-PoPS provides two complementary strategies to examine a prediction: 1) explanation by top contributor genes: Analogous to the representer values/points definitions in *Yeh et al.*^13^, K-PoPS reports the top n contributor (default n = 5) genes for each prediction ranked according to α*_i_k*(*x_t_*, *x_i_*). If these top contributor genes are functionally related to the trait, then the prediction is more likely to be biologically plausible. 2) explanation by anchor scores: K-PoPS allows users to pre-define a set of “anchor” genes that they believe are functionally linked to the trait *a priori*. These anchor genes could be curated from databases or literature, which represent known biological causes for a given trait. K-PoPS then aggregates the contribution scores from those anchor genes: 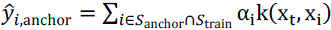, where *S*_anchor_ is the pre-defined anchor gene set. A low anchor score 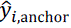 indicates the prioritized gene is not well-supported by the anchor genes, and therefore represent either unknown biology or potential a false positive prediction. Researchers can combine both explanations to evaluate a prediction: a prediction is more biologically plausible if its top contributor genes are closely related to the trait, and it has high anchor scores. Otherwise, the prediction is less trustworthy and should be treated with caution.

### Benchmarking identifies full-feature OLS ridge regression as the K-PoPS default

By default, PoPS first performs marginal feature selection using P value < 0.05, and then predicts MAGMA scores using generalized least squares (GLS) with ridge regularization. Marginal feature selection can substantially reduce the dimensionality of the feature matrix. This strategy is computationally attractive and was supported by the original PoPS parameter benchmarking, but its necessity is not guaranteed. Although the filtered features may be marginally uninformative, they might collectively show predictive power when modeled jointly. Additionally, ridge regression is designed to handle high degree of collinearity without requiring hard feature elimination^15–18^. Here, we first asked whether marginal feature selection is required for chromosome-held-out generalization in PoPS-style models, by benchmarking selected-feature versus full-feature implementations under otherwise comparable regression settings (selected features vs all). We additionally compared the ridge regression under ordinary least squares (OLS) and the GLS models (OLS vs GLS).

For benchmarking, we downloaded the summary statistics from Pan-UK Biobank (Pan-UKBB) for 38 traits, encompassing diverse trait categories such as metabolic, neurological, etc (Supplementary Table 1)^19^. These 38 traits represent a subset of the 113 traits analyzed by PoPS, and were selected based on our ability to match trait names between PoPS and Pan-UKBB summary statistics. We used the correlation between predicted and observed MAGMA scores as the evaluation metric 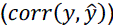. We found that OLS without feature selection yielded the best performance for 32/38 traits. GLS without feature selection yielded the best performance for 6/38 traits (Figure 2a and Supplementary Table 2). This observation suggests that marginal feature selection may discard predictive information and reduce generalizability. It also suggests that using OLS can slightly outperform the GLS alternative. Based on these initial benchmarking results, we implemented the dual form K-PoPS with OLS ridge regression without feature selection.

**Figure 2:**
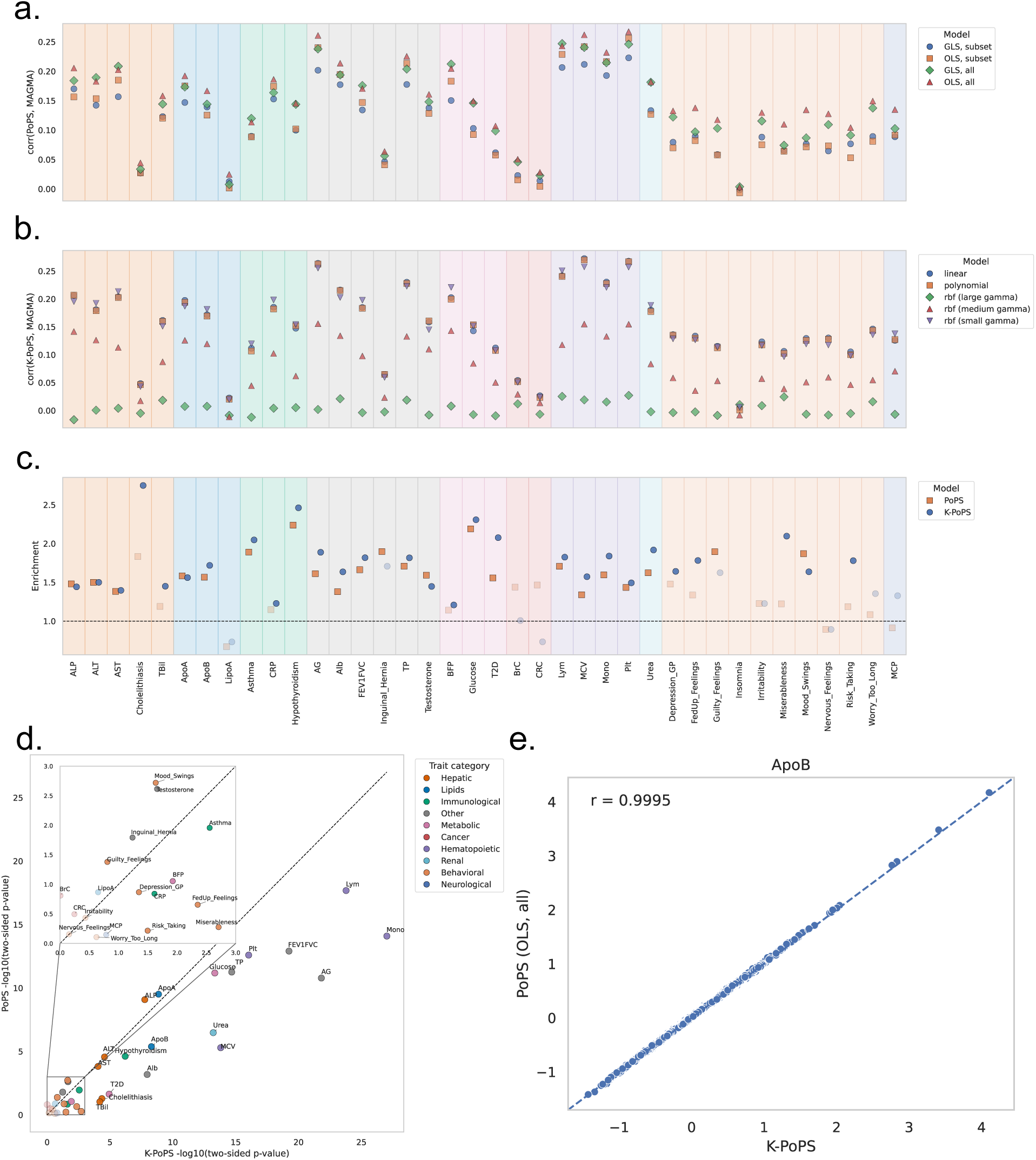
a) Comparing the predictive performance across 38 traits for 4 parameter settings with PoPS. The model using OLS and all features outperforms others for 32/38 traits. b) Five different kernel functions were benchmarked. The rbf kernel used hyperparameter γ = 1*e* − 6 (small), γ = 1*e* − 4 (medium), γ = 1*e* − 2 (large). We found the linear kernel, polynomial kernel (degree = 2), and rbf kernel (γ = 1*e* − 6) have comparable performance. c) Comparing the closest gene enrichment score between PoPS_GLS-selected_ and K-PoPS for 37 traits. Opaque data points are statistically significant (two side p < 0.05, calculated using matched moment method). Transparent data points are not statistically significant. Enrichment for “Insomnia” is not calculated since this trait has no GWAS hits. d) The enrichment P-value comparisons between PoPS and K-PoPS. e) The concordance between the non-default PoPS (OLS and all features) and K-PoPS for the ApoB trait.

The framework of K-PoPS enables the examination of other non-linear kernels. Here, we constructed the kernel matrix with other kernel functions, including polynomial and radial basis function (rbf) kernels. The comparison allows us to answer if non-linear functions can improve gene prioritization, which has not been systematically evaluated previously. Here, we constructed the kernel matrix with all the features, and compared the Pearson correlation between the LOCO-predicted and observed MAGMA scores. In general, we found that the linear kernel, polynomial kernel (degree = 2) and rbf kernel (γ = 1*e* − 6) can have very similar performance for predicting MAGMA on the unseen chromosome (Figure 2b and Supplementary Table 3). On the other hand, rbf kernels with larger gamma values can significantly underperform other models. This benchmarking experiment suggests that the linear kernel remains a robust choice for gene prioritization.

Following PoPS, we curated a dataset containing all the GWAS loci. The GWAS loci were defined as 1-Mb window around the lead SNP, which is obtained through LD clumping (Method). The gene prioritization scores were used to rank genes within a locus. Since ground-truth effector genes are generally unavailable, we used the closest gene to the lead SNP as a proxy ground truth for evaluation. Although not perfect, previous studies have shown that this strategy is enriched for true effector genes^20^, and this evaluation strategy was used in PoPS^6^. We were able to assign closest genes for 37 traits (Supplementary Table 4, “Insomnia” doesn’t have significant GWAS hits). By comparing K-PoPS with the default PoPS (PoPS_GLS-selected_), which employs GLS and feature selection step, we found that K-PoPS achieved higher enrichment score for 26/37 traits. Using p = 0.05 as a cutoff, K-PoPS score showed significant enrichment for 28/37 traits, while PoPS_GLS-selected_ score showed significant enrichment for 22 traits (Figure 2c, Supplementary Table 5). K-PoPS also tended to produce more significant p-values for the enrichment test, suggesting improved predictive power (Figure 2d). These comparisons indicate that the performance gain is primarily from using all features with OLS, whereas the kernel formulation provides the explanation framework. We demonstrate this by comparing K-PoPS and PoPS with an alternative formulation, which used the OLS with all the features (PoPS_OLS-all_). As expected from the equivalence between primal and dual ridge regression, K-PoPS produced highly concordant results with PoPS_OLS-all_(Figure 2e). Across all 38 traits, PoPS_OLS-all_ and K-PoPS scores had an average correlation = 0.9937 (Supplementary Table 6), consistent with the theory of kernel regression.

### Anchor scores stratify K-PoPS predictions by biological support

Since only a small fraction (0% – 33%) of PoPS prioritized genes have been functionally documented (Supplementary Table 7), we next ask whether gene prioritization could be improved by considering explanation evidence. We were able to define anchor gene sets for 25 out of 38 traits using orthogonal source of evidence. Among those 25 traits, we used HPO to define anchor gene sets for 7 disease traits, and used burden test results based on putative loss-of-function (pLoF) variants to define anchor gene sets for other 18 traits (Supplementary Table 1). We again defined GWAS loci based on independent lead SNP, and used K-PoPS to nominate genes within each locus.

We defined a “supported prediction” as a locus in which anchor scores and K-PoPS scores nominate the same gene, and an “unsupported prediction” as the locus in which anchor scores and K-PoPS scores nominate different genes. Treating the closest gene as the proxy ground truth, we defined a “correct prediction” for a locus if K-PoPS score nominated the closest gene. The locus is defined as an “incorrect prediction” if K-PoPS score nominated a different gene in a locus (Figure 3a). We used offset logistic regression to evaluate whether supported loci showed higher enrichment (for closest gene) than unsupported loci (Method). Offset logistic regression extends the matched moment framework introduced in PoPS^6^. This framework enables both estimation of baseline enrichment and direct comparison between supported and unsupported loci. When applied to baseline enrichment analysis, this statistical test is asymptotically equivalent to the matched moment approach proposed in PoPS^6^. We confirmed the consistency between the two statistical tests across all 25 traits. The resulting p-values were highly concordant. We also found that 21 of those 25 traits exhibits significant baseline enrichment (Figure 3b, Supplementary Table 8).

**Figure 3:**
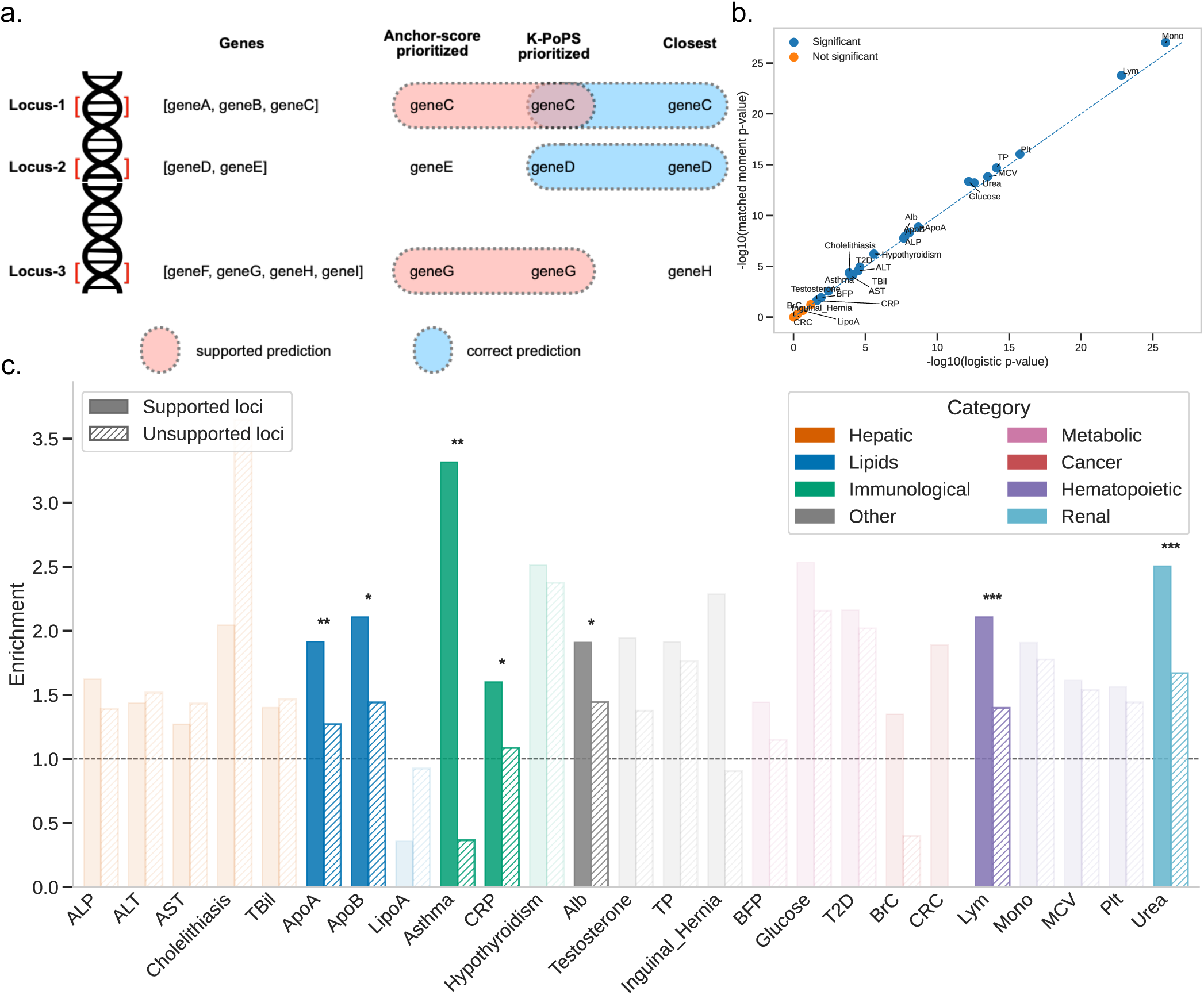
a) gene prioritization by locus. A correct prediction is the case when K-PoPS score prioritized the closest gene (deemed as the proxy effector gene). A “supported” prediction is the case when anchor scores and K-PoPS scores prioritize the same gene. b) Regardless of the anchor score, K-PoPS score alone have significant enrichment for 21 out of 25 traits. The x-axis is the p-value calculated with offset logistic regression, the y-axis is the p-value calculated with matched moment approximation, described in PoPS paper. Both methods computed highly consistent p-values. c) closest gene enrichment score for “supported” and “unsupported” predictions. Among 25 traits, 7 of them have statistically higher significant enrichment among supported predictions; 0 trait has significantly lower enrichment. p-value < 0.05 are labeled as *; p-value < 0.01 are labeled as **; p-value < 0.001 are labeled as ***.

We next applied offset logistic regression to compare supported and unsupported predictions. We reported two-side p-values for all 25 traits with offset logistic regression, and found that 7 traits showed differential enrichment between supported and unsupported predictions (Figure 3c, Supplementary Table 8). All these 7 traits have significantly higher enrichments among supported predictions, whereas no traits showed significantly lower enrichments. These results suggest that incorporating anchor scores improves effector gene (closest gene) nomination.

### Explainable prioritization for blood apolipoprotein B level with K-PoPS

Given that K-PoPS explanations improve effector gene prioritization, we further demonstrate its utility by examining a GWAS of blood apolipoprotein B (apoB) level. Pan-UKBB conducted GWAS on blood apoB level using 343,572 individuals^19^(phenocode: 30640). Elevated blood apoB levels are a well-known risk factor for cardiovascular disease^21^. We performed stringent LD clumping (see Method) on the summary statistics and identified 582 independent GWAS loci (Supplementary Table 4). We defined 18 anchor genes based on pLoF burden test results (Supplementary Table 1)^22^. We applied both PoPS and K-PoPS to nominate effector genes for each GWAS locus (Figure 4).

**Figure 4:**
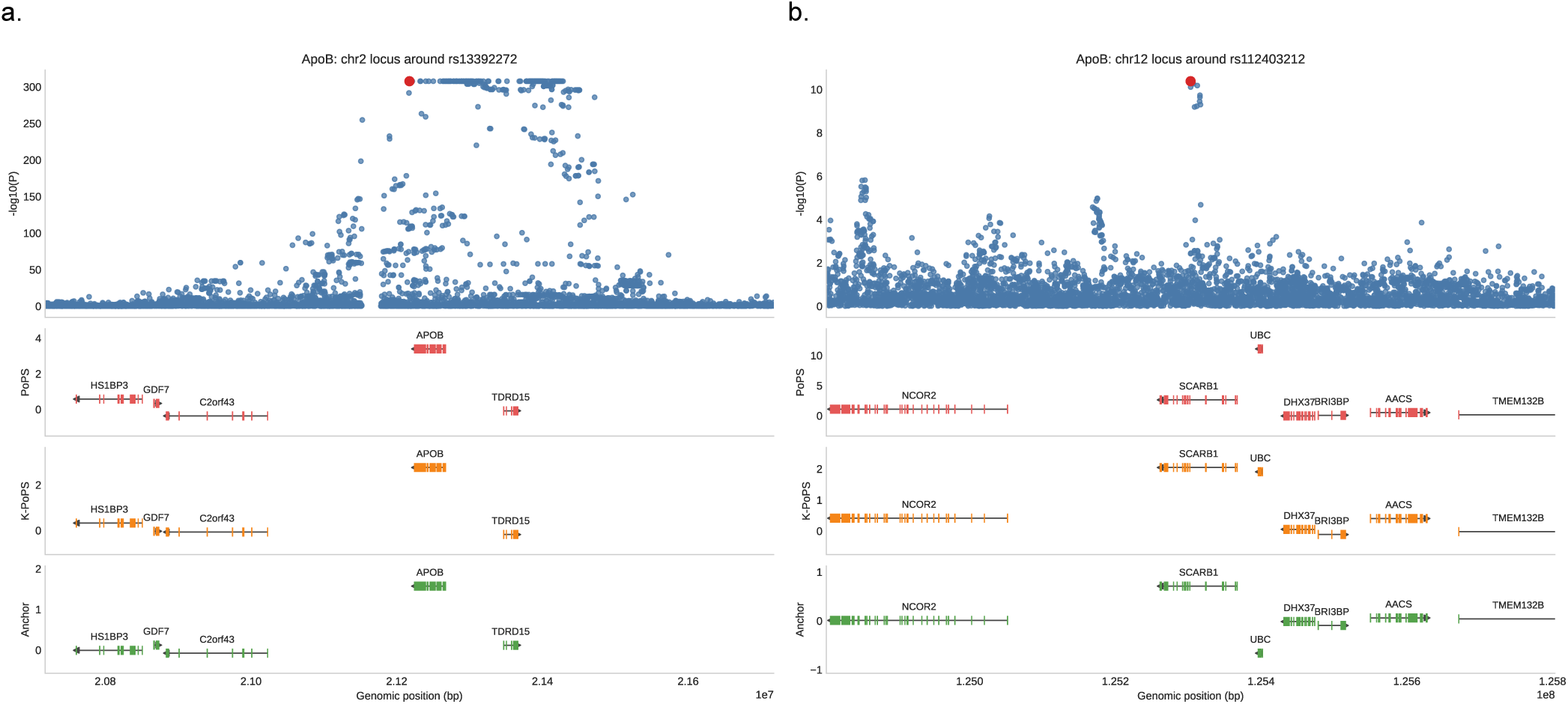
Examination of GWAS loci for blood apoB level. The top panel is a zoom-in Manhattan plot. The bottom 3 panels are prioritization scores for each gene in the locus. a) 1Mbps around rs13392272. Both PoPS and K-PoPS prioritized *APOB* as the effector gene. The prediction is supported by the anchor genes; b)1Mbps around rs112403212. PoPS prioritized *UBC*, while K-PoPS prioritized *SCARB1*. The anchor scores support *SCARB1* over *UBC*.

For the rs13392272 locus (chr2:20717490-21717490), both PoPS and K-PoPS identified the *APOB* as the effector gene (Figure 4a). This prediction is biologically plausible, since the *APOB* directly encodes apoB-48 and apoB-100, which are component of blood apolipoprotein B^21,23,24^. To explain the prediction, K-PoPS reported top 5 contributor genes for the APOB prediction, including *APOC3, APOA1, APOA4, APOA5, LPL*. These top contributor genes are generally in the same apolipoprotein family^25^, and encode proteins involved in lipid transport and metabolism, suggesting plausible prediction. An alternative explanation is to compare anchor scores within this locus. We found that *APOB* had the highest anchor score within this locus, suggesting that the prioritization of *APOB* is well-supported by pre-defined anchor genes (the bottom panel of Figure 4a). Taking both explanation evidence together, *APOB* is a biologically well-supported prediction by K-PoPS.

Next, we examined the rs112403212 locus (chr12:124803254-125803254). This locus contains *UBC*, which has the highest PoPS score genome-wide for blood apoB level (PoPS score = 11.08). The *UBC* gene encodes a component of ubiquitin complex and regulates numerous signaling pathways and cellular processes^26,27^. Functional studies have shown that ubiquitin is responsible for intracellular degradation of apoB protein^28^, although direct evidence linking *UBC* to circulating apoB levels is less established. On the other hand, a nearby gene, *SCARB1*, encodes plasma membrane receptor that transports high density lipoprotein cholesterol (HDL)^27,29,30^. PoPS prioritized *UBC* over *SCARB1*, while K-PoPS prioritized *SCARB1* over *UBC* (Figure 4b). Looking at the explanations, K-PoPS reported *SUMO1, PSMA5, TSG101, UBE2D3, TP53* as the top contributor genes for *UBC*. However, we found less trait-specific evidence connecting these contributor genes to circulating apoB levels, even though they show strong gene-level association signals (Supplementary Table 9). In contrast, the top 5 contributor genes for *SCARB1* are *LDLR, LPL, APOC3, APOA1, APOE*, all of which are well-established lipid genes^25^. Comparing anchor scores in this locus, we found that *SCARB1* is the highest while *UBC* is the lowest (bottom panel of Figure 4b). Combining both lines of explanation evidence, *SCARB1* is a more biologically plausible effector gene for blood apoB level in this locus.

### K-PoPS identifies plausible candidate genes in dilated cardiomyopathy locus

We next applied K-PoPS to dilated cardiomyopathy (DCM), a major cause of heart failure. DCM is estimated to have SNP heritability ranging from 11% – 20%, depending on the definition of the trait^31^. Zheng et al. performed comprehensive GWAS and post-GWAS analyses to identify putative effector genes. Using 8 different gene prioritization approaches (including PoPS), they prioritized 62 effector genes out of 80 genomic loci^31^. This study provides an ideal dataset to test K-PoPS, as it enables the evaluation of K-PoPS against other alternative gene prioritization approaches.

Because ground-truth effector genes are generally unknown, we used other seven methods (excluding PoPS, see Method) to define the putative ground-truth effector gene for each locus. Comparing PoPS (LOCO mode) with K-PoPS, we found that K-PoPS showed modestly higher enrichment for nominating putative ground-truth effector genes than PoPS (2.01 vs 1.75, Supplementary Table 10-11); but more importantly, it provided explanation evidence for locus-level interpretation. We defined anchor genes according to literature evidence (see Table 1 of *Eldemire et al.*^32^). *Eldemire et al.* further categorized these genes as “definitive”, “moderate”, “limited”. Using this categorization information, we were able to evaluate the precision-recall rate across anchor gene definitions for both supported and unsupported loci. We found that using all 51 anchor genes produced the best precision-recall rates for supported loci. Importantly, supported predictions exhibited substantially higher precision-recall rates than unsupported predictions (Figure 5a).

**Figure 5:**
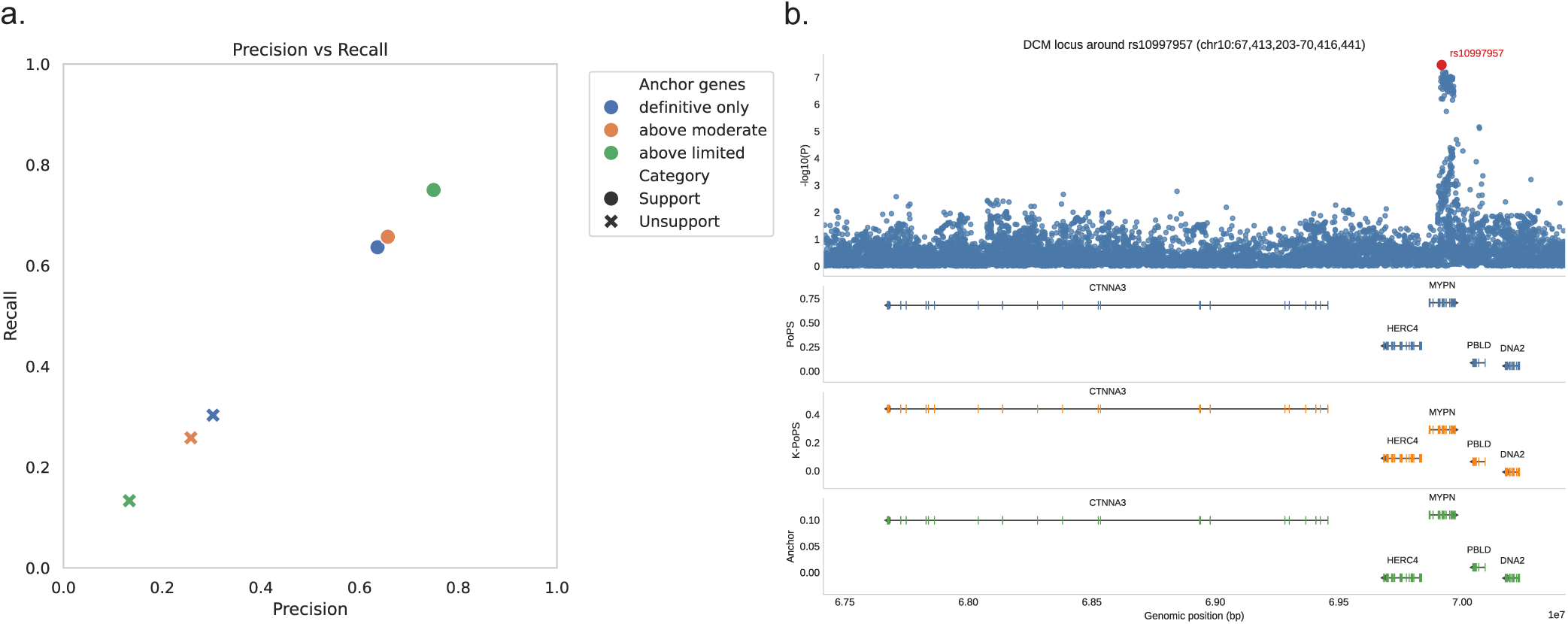
K-PoPS predictions on DCM. a) Precision-recall rate for supported vs unsupported loci across different anchor gene definitions. “definitive only” contains 12 anchor genes; “above moderate” contains 19 anchor genes; “above limited” contains all 51 anchor genes. b) Gene prioritizations for the locus around rs10997957. We extend the locus region (approximately 3Mbps) to include the gene body of *CTNNA3*. PoPS prioritized *MYPN*, while K-PoPS prioritized *CTNNA3*. *CTNNA3* and *MYPN* have comparable anchor scores.

Examining the predictions, we were surprised to find that K-PoPS prioritized *CTNNA3* over *MYPN* in this locus. The *MYPN* gene is the most likely effector gene in this locus, since six out of seven methods nominated this gene. This discrepancy between K-PoPS and other methodologies prompted us to further examine the model decisions. Looking at top contributor genes, we found that *CTNNA3* and *MYPN* have two shared top contributor genes: *ACTN2* and *HSPB7.* These top contributor genes have been reported to be involved in cardiomyopathy etiologies^33–35^ (Supplementary Table S9). We then compared the anchor scores in this locus, and we found that *CTNNA3* and *MYPN* exhibited similar anchor scores (Figure 5b. bottom panel). Combining both line of evidence, our analysis suggests that both *CTNNA3* and *MYPN* may contribute to DCM pathogenesis, challenging the common assumption that each GWAS locus contains a single effector gene. Indeed, *CTNNA3* has previously been implicated in DCM, and its phosphorylation is critical for maintaining cardiomyocyte-cardiomyocyte adhesion^36–38^. Notably, *CTNNA3* and *MYPN* have similar PoPS scores (Figure 5b. top panel), suggesting both genes are competitive candidates based on the prioritization score alone. However, K-PoPS provided an additional layer of explanatory evidence supporting this interpretation, offering further biological context for this hypothesis. In summary, incorporating explanatory evidence reveals that both *CTNNA3* and *MYPN* can be biologically plausible predictions.

## Discussion

Here, we present K-PoPS, a framework that enables explainable gene prioritization. Traditionally, similarity-based gene prioritization approaches assign a score to each gene, and nominate a single effector gene for each GWAS locus. This heuristic approach may nominate biologically implausible genes, and may also overlook genes worthy of further investigation. To address this problem, K-PoPS introduced post hoc explanations for each prediction. For a locus of interest, researchers can inspect if the prioritized genes are trustworthy, and if there are other biologically plausible candidates within the locus. K-PoPS achieves this goal by integrating the method of representer point into the PoPS framework^6,13^. By modeling all features with OLS, K-PoPS demonstrated improved predictive accuracy compared with default PoPS. Via kernel regression, K-PoPS transforms heterogeneous feature-centric explanations into intuitive gene-centric explanations. For blood apoB level, K-PoPS explanations provided supporting evidence for the prioritization of *APOB* gene, and supported prioritization of *SCARB1* over *UBC* as the putative effector gene. For DCM, K-PoPS revealed that both *CTNNA3* and *MYPN* may represent plausible effector genes, contrary to the one-gene-per-locus assumption. We hope this additional layer of information can guide biologists on decisions of downstream validations.

There are several notable limitations of K-PoPS: 1) quantitatively evaluating explanations is inherently challenging. Unlike supervised learning settings where ground truth labels are available, we generally don’t have well-defined metrics and ground truth for XAI tasks^39^. This is particularly true for gene prioritization tasks: the community has yet to define a widely accepted ground-truth effector gene list, let alone ground-truth model explanations. Nonetheless, we showed that supported predictions were more strongly enriched for putative ground-truth genes. 2) XAI approaches provide transparency regarding why a prediction is been made, but the explanations do not imply causation. Therefore, inferring mechanistic gene regulatory networks from contribution scores should be treated with caution; 3) K-PoPS is a reformulation of PoPS, and is model-specific. Compared to model-agnostic XAI approaches such as LIME^40^ and SHAP^11^, K-PoPS cannot be directly applied to other gene prioritization models such as DEPICT^5^. However, the concept of gene-centric explanation is extendible to more complex neural network models^13^. 4) We proposed defining the anchor genes *a priori* and aggregating their contribution scores. This approach is inherently sensitive to anchor-gene selection. Here, we defined anchor genes using orthogonal evidence derived from pLoF burden analyses^22^ and Human Phenotype Ontology (HPO) database^41^; 5) XAI provides guardrails for gene prioritization, which enables assessment of the credibility of each prediction. However, human judgement is still required for evaluating the biological plausibility of a prediction.

Numerous tools have been developed to prioritize genes for complex traits, but their explainability has been largely ignored. Here, we introduced K-PoPS, a similarity-based gene prioritization tool that enables interpretable predictions. We achieved this by attributing the prioritization score to the most influential training genes. Overall, we expect gene-centric explanations can lead to more transparent gene prioritizations, and facilitate downstream experimental decision-making.

## Method

### Pan-UKBB summary statistics

We downloaded GWAS summary statistics for 38 complex traits from the Pan-UKBB analysis conducted by the Neale lab (Supplementary Table 1)^19^. Pan-UKBB used approximately 330,000 individuals in its GWAS analyses, and included ‘*age, sex, age * sex, age*^2^*, age^2^ * sex, top 10 PCs*’ as covariates. Throughout this project, we used summary statistics derived from the European cohort. SNP coordinates from Pan-UKBB were based on the human genome Build 37 reference.

### MAGMA analysis

From the *MAGMA* website, we downloaded the Phase3 1000 Genomes reference panel (Build 37) of European ancestry. We downloaded the gene location file from *PoPS*, and reformatted it for compatibility with *MAGMA*. *MAGMA* annotated the SNP locations (in rsID) with the ‘--annotatè command. We then preprocessed the summary statistics by: 1) removing SNPs without rsIDs; and 2) SNPs with extremely small p-values (*p* < 2.2 × 10^−308^) were clipped to prevent numerical underflow. Following the recommendations of PoPS, we used the ‘snp-wise=mean’ for the MAGMA analysis^42^.

### PoPS analysis

We generally followed the PoPS GitHub workflow for this analysis^6^. Briefly, we downloaded the PoPS features from the Finucane lab. The gene feature file contains 57,742 heterogeneous features across 18,383 genes. PoPS first splits the feature file into NumPy chunks using ‘munge_feature_directory.py’. We then ran PoPS under a LOCO training scheme. Under the LOCO framework, we iterated over each chromosome and used only the training chromosomes for feature selection. The results from each chromosome were then concatenated. We reported the Pearson correlation between observed MAGMA score and PoPS score (LOCO) for each trait. We used the default parameter for PoPS, including: 1) regressing out MAGMA covariates; 2) using ridge regularization; 3) excluding the Human Leukocyte Antigen (HLA) region for training. In the first benchmarking experiment (OLS vs GLS, feature selection vs all features, Figure 2a), we re-implemented PoPS in PyTorch due to memory constraints (see Software implementation section). When comparing K-PoPS with the default PoPS model (GLS, feature selection) (Figure 2c), we used the original PoPS implementation. The re-implemented PoPS and the original PoPS produced very similar results (correlation > 0.99, Supplementary Table 6).

### K-PoPS analysis

We used the ‘prepare_kernel.py’ script to compute the kernel matrix using all PoPS features. We used the linear kernel throughout the analysis, but also benchmarked additional kernel functions (see Results section). The polynomial kernel was computed with degree = 2. We also computed the rbf kernels with the gamma parameter set to 1 × 10^−2^, 1 × 10^−4^, 1 × 10^−6^. Each feature was standardized to have mean 0 and standard deviation 1 prior to kernel calculation.

Using the kernel matrix, we then ran ‘k-pops.py’ with default parameters. In general, ‘k-pops.py’ behaves similarly to ‘pops.py’, except that it uses ridge-regularized OLS to predict MAGMA scores and uses all features for kernel construction. We used the LOCO framework and reported top five contributor genes for each prediction. We reported the Pearson correlation between observed MAGMA score and K-PoPS score (LOCO) for each trait.

### Anchor gene set curation

We defined anchor gene sets using orthogonal sources of evidence. Among the 38 traits, 7 of them (including CRC, BrC, Hypothyroidism, T2D, Asthma, Inguinal_Hernia, Cholelithiasis) had corresponding Phecode, which allowed us to map them to Human Phenotype Ontology (HPO)^41^. The Phecode-HPO map was curated by *McArthur et al*.^43^. The HPO database contains annotations of plausible effector genes for these traits from various sources. For the remaining 18 complex traits (mostly biomarkers and blood cell counts), we defined the anchor genes as genes harboring loss of function (pLOF) or missense variants, as reported by *Backman et al.* in the Whole Exome Sequencing (WES) analyses^22^ (Supplementary Table 1).

### Locus definition and gene prioritization criteria

We defined loci based on GWAS lead SNPs. We first performed stringent LD clumping using PLINK2^44^, with ‘--clump_r2 = 0.01, –-clump_kb = 500’, and restricted the analysis to genome-wide significant SNPs (*p* < 5 × 10^−8^). We then defined 1-Mb windows around lead SNPs and retrieved all genes within each window. Gene locations were defined as their Transcription Starting Site (TSS). To prioritize genes within a locus, we ranked all genes in the locus and selected the gene with the highest score. Because gene prioritization is not meaningful for loci without any genes, and is trivial for loci containing only one gene, we excluded these loci in the analysis. We also excluded loci with duplicated gene list, even if they were defined by different lead SNPs.

### Enrichment analysis for GWAS loci

For each locus, we ranked all genes by their K-PoPS scores and selected the gene with the highest score. Similarly, we selected the gene with the highest anchor score. If the anchor score and K-PoPS score selected the same gene within a locus, we call this a prediction with support from anchor genes (supported prediction); otherwise, we call this a prediction without support from anchor genes (unsupported prediction). We then treated the gene closest to the lead SNP as the putative effector gene. If K-PoPS selected the closest gene for that locus, we call this locus a correct prediction; otherwise, we call this an incorrect prediction.

Denote *x* as a binary variable indicating if the prediction of the locus *i* is supported by anchor genes (e.g. 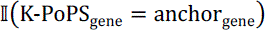, and *y* as a binary variable indicating the correctness of the locus 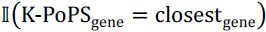. We fit a logistic regression with an offset: 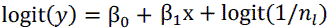. Here *n_l_* denotes the vector containing the number of genes in each locus. We used python package ‘statsmodels.GLM’ to calculate the p-value for β_1_. By default, ‘statsmodels.GLM’ applies a Wald test to calculate p-value. This offset logistic regression can also be used to evaluate whether prioritized genes show overall enrichment for the closest gene, by 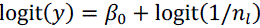. ‘statsmodels.GLM’ applies a Wald test to test if β_0_ is significantly deviating from 0. Empirically, this statistical test was consistent with the two-sided p-values calculated using 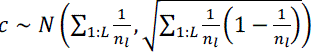, as proposed by PoPS^6^. We used the matched moment approach to calculate the enrichment shown in Figure 2c.

### Gene prioritization for DCM

Following the analysis by *Zheng et al.*, we used meta-analyzed GWAS summary statistics defined using broad diagnostic criteria, comprising 14,256 cases and 1,199,156 controls^31^. Anchor genes were defined based on Table 1 from *Eldemire et al.*, which we further categorized into “definitive only” (12 genes), “above moderate” (19 genes), and “above limited” (51 genes)^32^. We ran K-PoPS under the LOCO setting and compared their precision and recall across different definitions for anchor genes. Putative ground-truth effector genes were defined using the total score from seven methods including “ABC”, “TWAS”, “Colocalisation”, “V2G”, “Coding variant”, “Nearest”, “Mendelian disease-causing gene” (see Supplementary Table 8 of *Zheng et al.*). Among these 80 loci, we were able to assign putative ground-truth effector genes for 66 loci, comprising 309 genes in total. All these loci were partitioned into “supported” and “unsupported”, based on the agreement between the K-PoPS scores and anchor scores. We then evaluate the precision and recall rate for these predictions. We followed PoPS to define True Positives (TP), True Negatives (TN), False Positives (FP), and False Negatives (FN), and to calculate precision and recall^6^. Because an incorrect prediction can simultaneously create one FP and one FN, the resulting precision was always equal to recall.

### Software implementation

To enable benchmarking using the full feature set, we re-implemented PoPS in PyTorch using lower-precision floating point arithmetic (float32). This new implementation was more memory efficient and enabled prediction using more than 55,000 features. We also optimized the LOCO scheme so that the MAGMA gene-level covariance matrix was loaded only once, thereby improving computational efficiency.

Similarly, we implemented K-PoPS with PyTorch using float32. K-PoPS natively supports LOCO computation. It generally follows PoPS, except that it can report the top n contributor genes (from the training chromosome) for each prediction. It can also compute the contribution scores for the anchor genes (also from the training chromosome). Similar to PoPS, the optimal regularization parameter was determined using Leave-One-Out-Cross-Validation (LOOCV) among the training genes. The PyTorch implementation allows GPU computation, which improved computational speed by approximately 20-fold (Supplementary Table 12).

## Data & Code availability

Pan-UKBB summary statistics: https://pan.ukbb.broadinstitute.org/

MAGMA (tool and reference panel): https://cncr.nl/research/magma/

PoPS features: https://www.finucanelab.org/data

PoPS: https://github.com/FinucaneLab/pops/tree/master

Manuscript Pipeline: https://github.com/Samee-Lab/k-pops-manuscript

K-PoPS: https://github.com/JasonTan-code/k-pops

DCM summary statistics: https://www.nature.com/articles/s41588-024-01952-y

DCM anchor genes: https://pmc.ncbi.nlm.nih.gov/articles/PMC10842880/

## Acknowledgement

We thank Dr. Katherine Pollard for her invaluable feedback to this project. This project is supported by R01HL175964, R01HL169511 and R01CA288967.

## Author contributions

T.T. and M.A.H.S wrote and reviewed the manuscript. T.T. conceived, implemented the model, and conducted data analysis. M.A.H.S advised on the project.

